# Characterization of *SETD1A* haploinsufficiency in humans and *Drosophila* defines a novel neurodevelopmental syndrome

**DOI:** 10.1101/2019.12.17.879189

**Authors:** Joost Kummeling, Diante E Stremmelaar, Nicholas Raun, Margot RF Reijnders, Marjolein H Willemsen, Martina Ruiterkamp-Versteeg, Marga Schepens, Calvin CO Man, Christian Gilissen, Megan T Cho, Kirsty McWalter, Margje Sinnema, James W Wheless, Marleen EH Simon, Casie A Genetti, Alicia M Casey, Paulien A Terhal, Jasper J van der Smagt, Koen L van Gassen, Pascal Joset, Angela Bahr, Katharina Steindl, Anita Rauch, Elmar Keller, Annick Raas-Rothschild, David A Koolen, Pankaj B Agrawal, Trevor L Hoffman, Nina N Powell-Hamilton, Isabelle Thiffault, Kendra Engleman, Dihong Zhou, Olaf Bodamer, Julia Hoefele, Korbinian M Riedhammer, Eva MC Schwaibold, Velibor Tasic, Dirk Schubert, Deniz Top, Rolph Pfundt, Martin R Higgs, Jamie M Kramer, Tjitske Kleefstra

## Abstract

Defects in histone methyltransferases (HMTs) are major contributing factors in neurodevelopmental disorders (NDDs). Heterozygous variants of *SETD1A* involved in histone H3 lysine 4 (H3K4) methylation were previously identified in individuals with schizophrenia. Here, we define the clinical features of the Mendelian syndrome associated with haploinsufficiency of *SETD1A* by investigating 15 predominantly pediatric individuals who all have *de novo SETD1A* variants. These individuals present with a core set of symptoms comprising global developmental delay and/or intellectual disability, subtle facial dysmorphisms, behavioral and psychiatric problems. We examined cellular phenotypes in three patient derived lymphoblastoid cell lines with three variants: p.Gly535Alafs*12, c.4582-2_4582delAG, and p.Tyr1499Asp. These patient cell lines displayed DNA damage repair defects that were comparable to previously observed RNAi-mediated depletion of *SETD1A*. This suggested that these variants, including the p.Tyr1499Asp in the catalytic SET domain, behave as Loss-of-Function (LoF) alleles. Previous studies demonstrated a role for SETD1A in cell cycle control and differentiation. However, individuals with *SETD1A* variants do not show major structural brain defects or severe microcephaly, suggesting that defective proliferation and differentiation of neural progenitors is unlikely the single underlying cause of the disorder. We show here that the *Drosophila Melanogaster* SETD1A orthologue is required in postmitotic neurons of the fly brain for normal memory, suggesting a role in post development neuronal function. Together, this study defines a neurodevelopmental disorder caused by dominant *de novo* LoF variants in *SETD1A* and further supports a role for H3K4 methyltransferases in the regulation of neuronal processes underlying normal cognitive functioning.

## Introduction

Neurodevelopmental disorders (NDDs) comprise a heterogeneous group of disorders including intellectual disability (ID) and autism spectrum disorders (ASDs)^1^. Approximately 1:200 to 1:450 children are born with a *de novo* pathogenic gene variant causing a Mendelian NDD^2^. Interestingly, recent large-scale exome sequencing studies have shown that defects in histone methyltransferases (HMTs) are major contributors to ASD and ID^3, 4^.

One of the genes involved in epigenetic modification through histone methylation is *SETD1A* (also known as *KMT2F*), which encodes a histone methyltransferase that mediates trimethylation of histone H3 on lysine 4 (H3K4me3) at promoters of active genes^5^. SETD1A methyltransferase activity depends on interactions with other proteins in a highly conserved complex, called ‘complex of proteins associated with Set1’ (COMPASS). COMPASS was originally defined in yeast^6^, and subsequently shown to be conserved in flies and mammals^7–9^.

*SETD1A* appears to be important for multiple aspects of cell cycle regulation. For example, SETD1-mediated H3K4me3 activates β-catenin expression, which is required for proliferation of neuronal progenitor cells^10^. Furthermore, *SETD1A* has been shown to promote cell cycle progression by activating the expression of micro RNAs that suppress the antiproliferative gene *BTG2*, which plays a role in the transition between the G1 and S phase of the cell cycle^11^. Finally, H3K4 methylation by *SETD1A* is crucial in maintaining genome stability, especially during DNA replication, by protecting newly-replicated DNA from degradation^12^.

Heterozygous Loss-of-Function (LoF) variants of *SETD1A* have been identified in several human individuals from large schizophrenia cohort studies and have been linked to disruption of speech development and early-onset epilepsy^13–15^. Previously, LoF variants were also identified in a limited number of children with developmental delay^14^, suggesting that *SETD1A* variants are a common biological underpinning of the different aforementioned phenotypes. To provide further insight into the somatic and behavioral profiles, we characterised the neurodevelopmental syndrome associated with *SETD1A* heterozygous LoF variants in this study. This is a prerequisite to develop future guidelines and management directed towards the vulnerability to schizophrenia or other psychiatric diseases. In addition, to further establish the role of *SETD1A* in underlying biological processes, we studied the effect of *SETD1A* variants on genome stability in patient cell lines, and investigated the role of the *Drosophila Melanogaster* orthologue, Set1, in postmitotic memory forming neurons of the fly brain.

## Methods

### Patients

We collected the molecular and clinical features of 15 unpublished individuals with *de novo SETD1A* variants by a collaboration facilitated by GeneMatcher^16^ in which several clinical groups independently identified individuals with developmental delay/intellectual disability (DD/ID) and related phenotypes with rare variants in *SETD1A* during routine diagnostic genetic testing. Clinical analysis of these patients was performed during regular consultations focusing on medical history, physical examination, and observational analysis of behavioral features. In all patients, exome sequencing and variant filtering were performed, according to the routine procedures at each institute^17–22^.

We compared these data to available information from previously published cases with pathogenic *SETD1A* variants associated with schizophrenia and NDDs^14, 23, 24^. Ten individuals reported in these previous studies originate from cohorts diagnosed with schizophrenia (UK10K-Finns and Swedish cohorts) and six individuals originate from cohorts of individuals seen with NDDs (Deciphering Developmental Disorders (DDD) project, Northern Finland Intellectual Disability (NFID) study and Northern Finland Birth Cohorts (NFBC)). Clinical information of the previously reported individuals was only scarcely provided.

To further assess the impact of these variants on cellular SETD1A functions, we first isolated and generated lymphoblastoid cell lines (LCL) from 3 patients (6, 13 and 15), and from two unrelated controls.

For further details on the methods used for cell culture, immunoblotting, antibodies and RNA isolation see Supplementary Data 1 - Methods.

### DNA Fibres

DNA fibre analysis was carried out as described previously^12^. Briefly, cells were pulse-labelled with CldU and IdU (Sigma-Aldrich) for 20 min each before a 5 h exposure to 4 mM HU, and at least 200 replication forks were analysed per experimental condition. Images were taken using a Nikon Eclipse Ni microscope equipped with a 60X oil lens, and were acquired and analysed using Elements v4.5 software (Nikon). The lengths of labelled tracts were measured using ImageJ^25^ and arbitrary length values were converted into micrometers using the scale bars created by the microscope.

### Drosophila Stocks and Genetics

To study the role of Set1 in postmitotic memory forming neurons of the fly brain, flies were reared on a standard medium (cornmeal-sugar-yeast-agar) at 25°C and 70% humidity with a 12 h: 12 h light/dark cycle. *R14H06-Gal4* flies express Gal4 under the control of a mushroom body (MB) specific enhancer from the *rutabaga* gene^26, 27^. *mCherry^RNAi^*, *Set1^RNAi1^*, *Set1^RNAi2^*, and *Set1^RNAi3^* were all generated by the Transgenic RNAi Project at Harvard University Medical School^28^ using a common donor strain, therefore these RNAi lines all share a common genetic background. *mCherry^RNAi^* was used as a control in these experiments because it shares a common genetic background with all of the tested RNAi lines, and represents any non-specific effects of a non-targeting RNAi. *mCherry^RNAi^*, *Set1^RNAi1^*, *Set1^RNAi2^*, and *Set1^RNAi3^* were crossed to *R14H06-Gal4* for courtship conditioning and activity experiments, *actin-Gal4* for lethality assays, and *UAS-mCD8::GFP; R14H06-Gal4* for analysis of MB morphology. For more details on the acquisition of the used fly stocks, see Supplementary Data 1 - Methods.

### Courtship Conditioning

Flies were trained and tested for short-term (STM) and long-term memory (LTM) 4 days after eclosion using the courtship conditioning assay as previously described^29–31^. For each fly pair a courtship index (CI) was calculated, which is the proportion of time spent courting over 10 minutes. The learning Index (LI) represents the percentage of reduction in courtship behavior due to training and is used to directly compare memory ability between different genotypes. LI is a single value that is calculated from the CIs using the formula: LI= (mean-CInaïve – mean-CItrained)/ mean-CInaïve. For more details on the courtship conditioning procedures, see Supplementary Data 1 - Methods.

### Analysis of Activity and Sleep

One-to five-day old flies were assayed for locomotor activity in 12 h: 12 h light/dark regimens after at least three days entrainment, for four full days using DAM5M monitors (Trikinetics, Waltham, MA). Locomotor data was collected in 1 minute bins, and a 5 minute period of inactivity was used to define sleep^32, 33^. Sleep parameters were analysed using pySolo and plotted^34^. Dead animals were excluded from analysis by a combination of automated filtering and visual inspection of locomotor traces.

### Analysis of Mushroom Body Morphology

Brains from one-to five-day old day old adult male flies expressing the membrane bound UAS-mCD8::GFP under the control of R14H06-Gal4 were dissected in 1x PBS. Dissected brains were fixed in 4% paraformaldehyde for 50 minutes and then washed in a PBS with 1% Triton X-100 for 5 minutes. Brains were then mounted using Vectashield and imaged with a Zeiss LSM 710 confocal microscope. Confocal projections were created using ImageJ software (v. 2.0.0)^25^.

## Results

### Patients: Molecular Phenotype

An extensive overview of the clinical and molecular data of the patients included in this study is provided per individual in Supplementary Table 1. For one variant paternal genotype data was unavailable, therefore paternal inheritance could not be fully excluded. All other variants of the *SETD1A* gene identified in our cohort had occurred *de novo.* The 15 *SETD1A* variants identified here include 5 nonsense, 6 frameshifts, 1 missense and 2 different splice site variants, of which one occurred in two unrelated individuals. These 14 different identified *SETD1A* variants were all located before the conserved SET domain, which is responsible for catalyzing H3K4 methylation (Figure 1).

**Figure 1.**
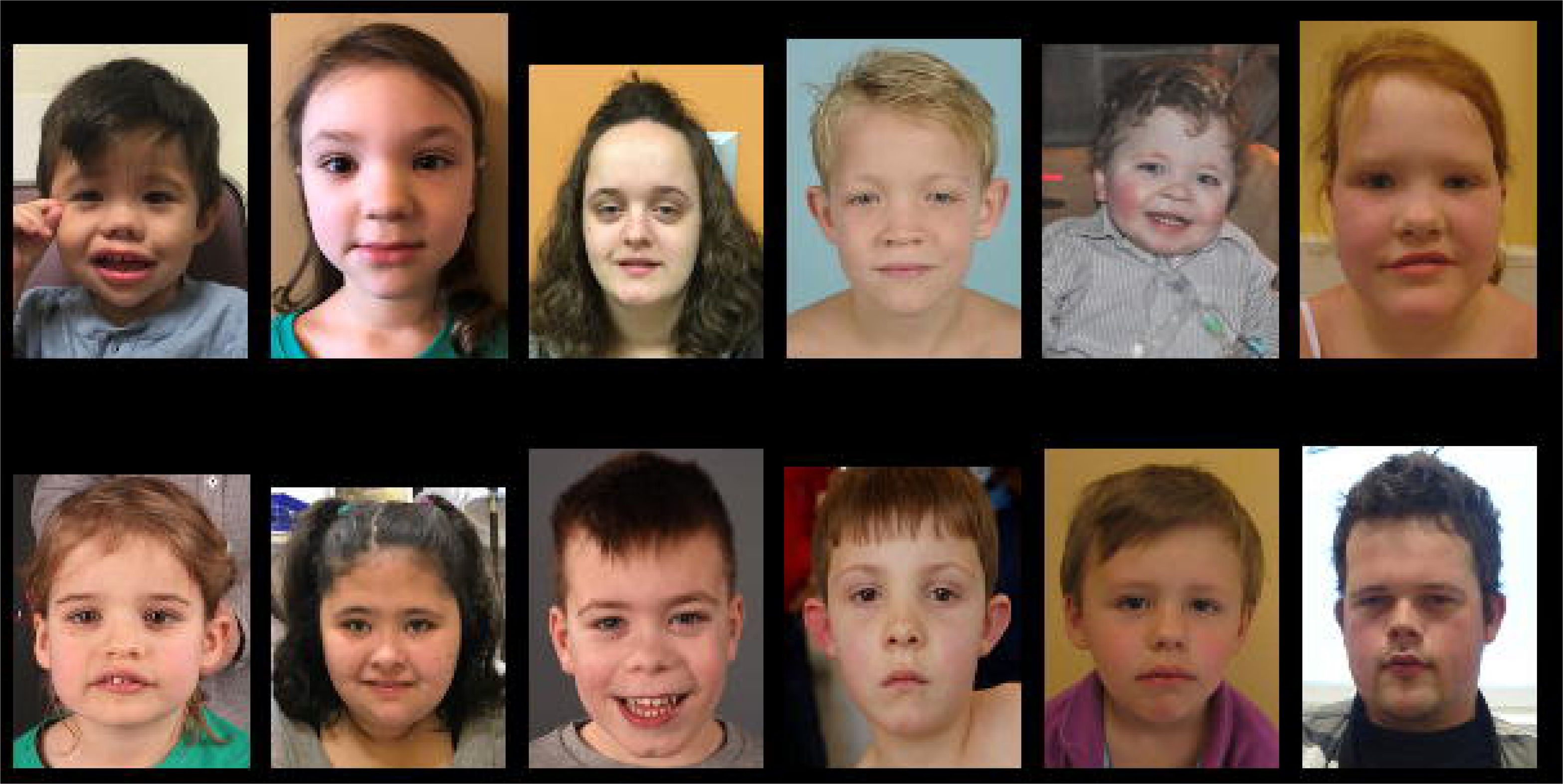
The genomic position and coding consequences of 14 different *SETD1A* variants. All variants are located 5’ to the conserved SET domain, which is responsible for catalyzing methylation. The splice acceptor variant at exon 16 (c.4582-2delAG) was identified in two unrelated patients. Mapping of the different domains (IPR024657, IPR001214, IPR000504 and IPR003616) on the gene was done according to data retrieved from Interpro^61^.

To gain further insight into the pathogenicity of the missense variant (Tyr1499Asp), we utilized several *in silico* prediction models. MetaDome is an online data platform that uses homologous domain relations and combines data from GnomAD with ClinVar to produce a ‘tolerance landscape’ of a gene^35, 36^. According to MetaDome the region in which the variant lies is intolerant (Supplementary Figure 1). In addition, several other *in silico* prediction tools indicated that this variant is pathogenic: CADD PHRED: 29.7^37^; SIFT: Deleterious, score 0^38^; Polyphen-2: Probably damaging, score 1, 000^39^.

In each patient, the observed *SETD1A* variant was considered to be the most likely contributor to the phenotype. In general, *SETD1A* is depleted from truncating variants in healthy individuals (pLi=1). Only 2 of the 14 different variants were listed in GnomAD: c.2968C>T and c.4582-2_4582delAG (with a low allele frequency of 4.061e^-6^ and 8.237e^-6^, respectively)^40^. Interestingly, this latter variant, (c.4582-2_4582delAG) was identified in two different patients but had also been identified previously^14^. To determine any effects of this splice-site variant on the *SETD1A* transcript, RT-PCR was performed on cells from individual 15, amplifying a region between exon 14 and 18. The resulting product shows that the transcript produced in individual 15 differs from the wildtype transcript. Sequencing of the amplified PCR-products revealed that the variant causes an insertion of 81 base pairs between exon 15 and 16, showing that intron 15 is not spliced. This insertion is in frame, but the N-SET domain, which is important for methyltransferase activity, is interrupted (Supplementary Figure 2). Preventing nonsense mediated RNA decay with cyclohexamide resulted in a 50:50 ratio of wildtype (WT) to mutant transcript. Without cyclohexamide the WT product was higher than mutant, suggesting that the mutant mRNA is partly degraded in the cell. These results suggest that this recurrent splice acceptor variant causes LoF by disrupting the N-SET domain of SETD1A.

### The Cellular Impact of SETD1A Variants

Immunoblotting analyses revealed that cells from two patients, harboring the p.Gly535Alafs* and c.4582-2_4582delAG variants, exhibited decreased levels of SETD1A protein. The presence of the Tyr1499Asp variant had no effect on SETD1A protein levels (Figure 2A). Moreover, when these patient-derived cells were exposed to DNA damage in the form of high-dose hydroxyurea, they exhibited slightly elevated activation of the DNA damage response compared to WT counterparts, evidenced by increases in phosphorylation of CHK1 and RPA (Figure 2B).

**Figure 2.**
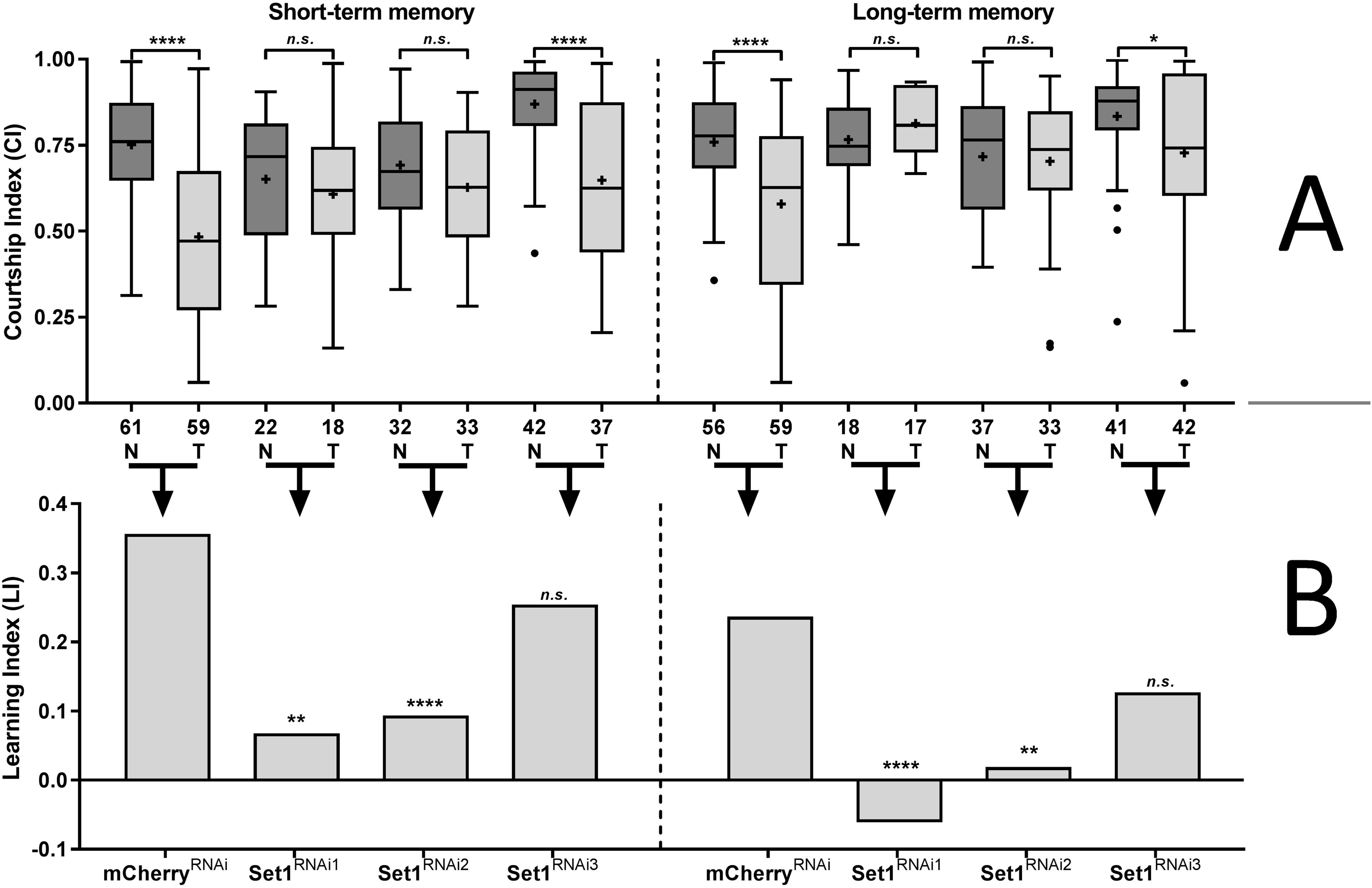
(A) Whole cell extracts (WCE) from the indicated patient-derived lymphoblastoid cell lines (LCL) were analysed by immunoblotting using the denoted antibodies. Data is derived from the untreated samples in (B). (B) Patient-derived LCL were exposed to 2 mM HU for 24 h, WCE prepared and analysed by immunoblotting using the denoted antibodies. Data is representative of n=2 independent experiments. * Denotes a non-specific band. (C) Patient-derived LCL were sequentially pulsed for 20 min each with CldU and IdU, and exposed to 4 mM HU for 5 h (as in the schema). DNA was visualised with antibodies to CldU and IdU, and tract length was calculated. Plots denote the average ratios of IdU:CldU label length from three independent experiments. Red lines indicate mean ratios. Data represents pooled values from n=3 experiments, representative images are shown.

More strikingly, *SETD1A* variants were also associated with severe levels of nascent DNA degradation upon DNA damage, to levels comparable with depletion of SETD1A^12^. This was not evident in control cells, but was apparent in all 3 patient-derived cell lines examined (Figure 2C). These data suggest that these variants, including the Tyr1499Asp variant within the catalytic SET domain, may all lead to loss of SETD1A function. Moreover, given the severity of this defect, heterozygous variants in *SETD1A* mimic phenotypes that were previously observed in cell lines treated with SETD1A siRNA.

### Patients: Clinical Phenotype

Our cohort of 15 individuals with heterozygous *SETD1A* variants comprised 8 females and 7 males whose ages varied from 34 months to 23 years (Supplementary Figure 3). The collection of clinical data of these individuals has allowed us to further define the clinical phenotype associated with *SETD1A* haploinsufficiency. A summary of the clinical data of the 15 patients included in this study is presented in Table 1. Figure 3 depicts the facial appearance of the patients for which portrait photographs were available. An extensive overview of the clinical features of the 15 individuals can be found in Supplementary Table 1. For comparison, we provide a synopsis of the clinical descriptions of the previously published cohort of 16 individuals with *SETD1A* variants^14^ in addition to our clinical data (Column 17 in Supplementary Table 1). For additional detailed clinical information per individual, see Supplementary Data 2 - Results.

**Figure 3.**
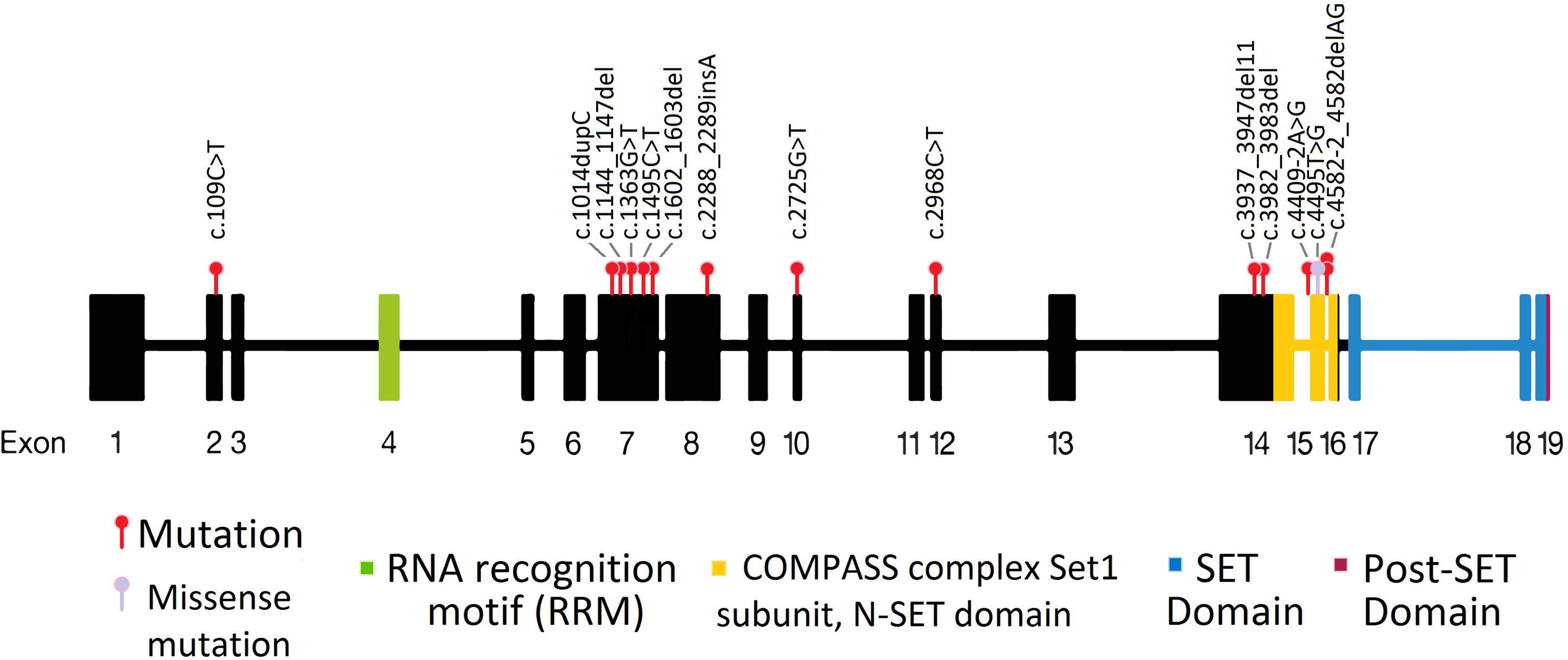
Clinical portrait photographs of patients with a *SETD1A* pathogenic variant. The most prominent facial features to be observed are high forehead (7/15), low-set ears (4/15), microtia (3/15), downslanted palpebral fissures (6/15), epicanthus (7/15), deeply set eyes (4/15), hypertelorism (4/15), wide nose (6/15), anteverted nares (4/15), full cheeks (2/15), everted/tented upper lip vermillion (8/15), wide mouth (4/15) and widely spaced teeth (3/15).

**Table 1.**
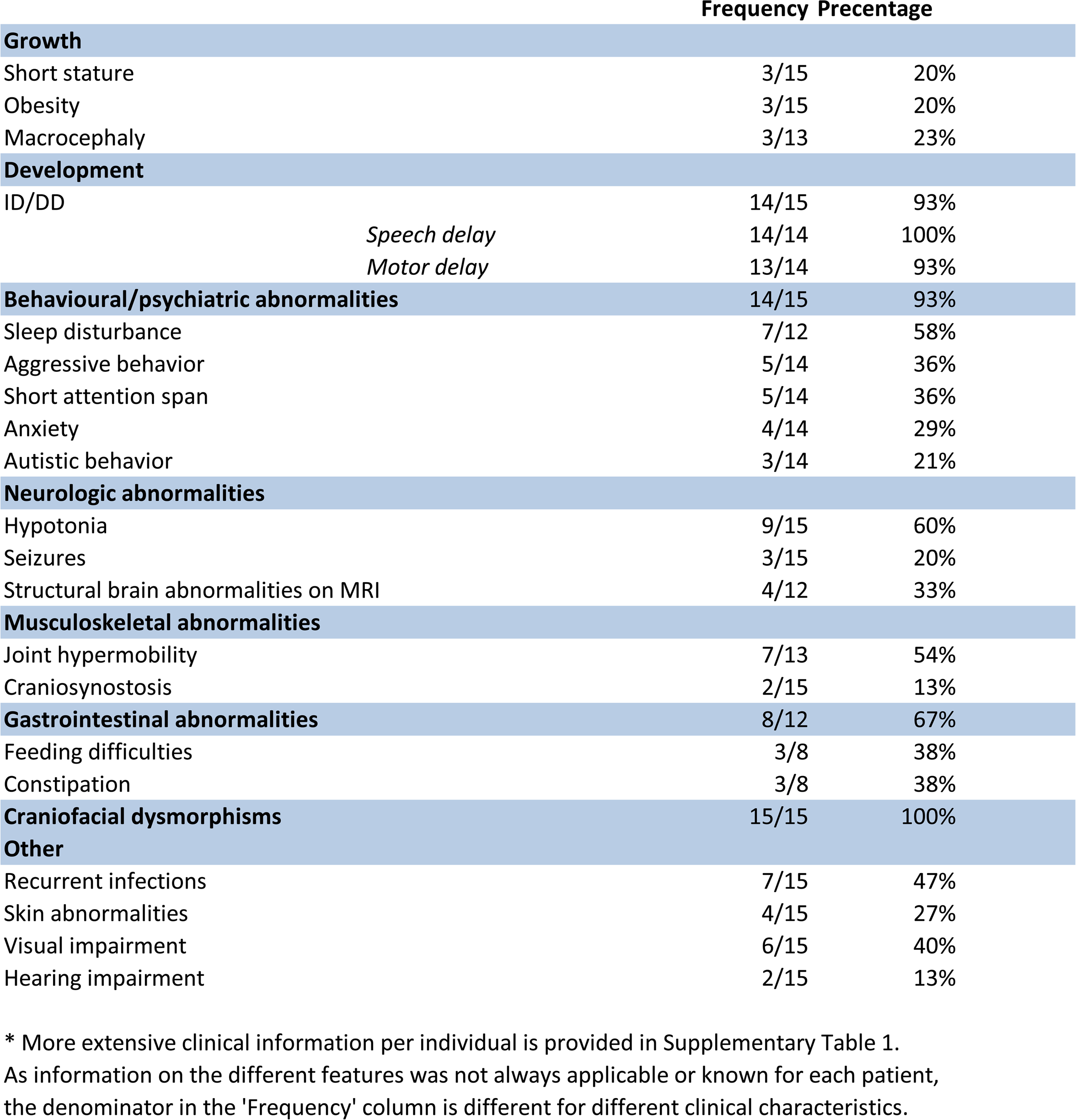
Summary of clinical features found in the cohort of 15 individuals with SETD1A variants.* * More extensive clinical information per individual is provided in Supplementary Table 1. As information on the different features was not always applicable or known for each patient, the denominator in the ’Frequency’ column is different for different clinical characteristics.

### Development

Global developmental delay was reported in 14/15 individuals, including delayed speech and language development (14/14) and motor development (13/14). In 6 patients formal IQ evaluations have been performed, confirming intellectual disability (TIQ < 70) in 5/6 patients. In the other individuals, observations in educational environments indicated learning difficulties or developmental delay in 6/9 individuals. The level of ID or developmental delay is generally mild.

### Behavioral and Psychiatric Abnormalities

Because of the previously reported association between schizophrenia and variants in the *SETD1A* gene^14^, we specifically looked at psychiatric and behavioral abnormalities in our patient cohort. Behavioral abnormalities were observed in 14/15 individuals. These consisted of aggressive behavior (5/14), anxiety (4/14), short attention span (4/14), autistic features (3/14), overfriendliness (3/14) and abnormal temper tantrums (2/14). In our cohort, 7 out of 12 patients suffered from sleep disturbances, including night terrors.

There were signs (though no formal diagnoses) of a psychotic disorder in two individuals in the cohort. First, case 15, a 23-year-old man with the recurrent splice acceptor change (c.4582-2_4582delAG), experienced periods with apparent negative symptoms associated with schizophrenia (somberness and social withdrawal), alternating with periods of euphoria. He was treated with aripiprazole, 10 mg daily. He also showed aggressive and addictive behavior from a young age and has been diagnosed with PDD-NOS (DSM-IV criteria), which is part of the autism spectrum according to current diagnostic criteria (DSM-V)^1^. Second, case 3, a 16-year-old female, showed symptoms that could be classified as visual hallucinations and imperative auditory hallucinations, as she was reported to “hear voices telling her to do things and see things that are not there”. This seemed to be transient and has not reoccurred. Because she was assessed as doing well, she so far has received neither formal evaluation nor medication for these symptoms.

### Neurological Abnormalities

Seizures were present in 3/15 individuals, with some very severe forms. For example, case 2 had myoclonic-astatic epilepsy. Drug treatment was sufficient to control seizures in case 2 and 3, but for case 11 hemispherectomy was required for seizure control. Hypotonia was reported in 9/15 individuals, often manifesting during infancy and remained relatively mild. In 4/12 patients for which MRI has been performed, morphological abnormalities of the brain were reported including corpus callosum abnormalities, cerebral white matter abnormalities, T2 hyperintensities, and Chiari I malformation. Other reported neurological abnormalities are EEG abnormalities, encephalopathy, dysarthria, slight tremor, sensory processing disorder, and short-term memory deficit.

### Musculoskeletal Abnormalities

Musculoskeletal abnormalities mainly affect the extremities (12/15) and included pes planus (6/12), broad/wide fingers (3/12), long fingers (2/12), and tapered fingers (2/12). Joint hypermobility was reported for 7/13 individuals. Other, less penetrant musculoskeletal abnormalities included plagiocephaly (2/15), hyperlordosis (2/15), craniosynostosis (2/15), kyphosis (1/15), pectus excavatum (1/15), hip dysplasia (1/15), postural instability (1/15), right sided preference (1/15), and congenital torticollis (1/15).

### Gastrointestinal Abnormalities

Gastrointestinal abnormalities were present in 8/12 individuals. Feeding difficulties were reported for 3/8 individuals. The feeding difficulties mainly occurred and resolved during infancy. Constipation was observed in 3/8 individuals. Less common features included gastroesophageal reflux, frequent bowel movements and insatiable appetite.

For additional clinical information on highlighted individual cases see Supplementary Data 2 - Results.

### Drosophila: Memory Phenotype

To gain insight into the function of SETD1A in neurons, we investigated *Drosophila* Set1, which is an orthologue of human SETD1A and SETD1B. Both *SETD1A* and *SETD1B* are mutated in individuals with overlapping neurodevelopmental disorders^41^. The Set1 family of proteins are known to be important regulators of cell proliferation and differentiation^10–12, 42, 43^. However, phenotypes that are normally associated with altered cell proliferation, *i.e.* overgrowth and cancer, are not frequently observed in *SETD1A* or *SETD1B* patient cohorts. To gain insight into the role of *Drosophila* Set1 in other aspects of neurobiology, we investigated terminally differentiated postmitotic neurons of the *Drosophila* mushroom body (MB), the primary learning and memory center of the fly brain.

Since germline Set1 mutants are lethal^44^, we used the spatially restricted Gal4/UAS system^45^ to induce targeted RNA interference (RNAi) of Set1. Postmitotic MB neurons were targeted using the *R14H06-Gal4* driver line, which we have characterised previously^26, 27^. *R14H06-Gal4* is highly specific for the MB in adult and larval fly brains^46^. It is also expressed in a few cells of the ventral nerve chord, which appear to be chordotonal sensory neurons involved in balance and locomotion. We obtained three different Set1 UAS-RNAi transgenes that target different parts of the Set1 mRNA. We tested the RNAi potency by crossing RNAi stocks to the ubiquitous actin-Gal4 driver. As Set1 is an essential gene this would be expected to induce lethality. While *Set1^RNAi1^* and *Set1^RNAi2^* did induce lethality, *Set1^RNAi3^* did not, suggesting that the *Set1^RNAi3^* line is less potent. As a control, we used an mCherry RNAi strain (*mCherry^RNAi^*) that had the same genetic background as the Set1 RNAi strains.

We tested the effect of postmitotic Set1 knockdown on memory using the courtship conditioning paradigm, a classic memory assay based on naturally occurring courtship behaviors^29^. Socially naïve male flies court female flies vigorously, however previously mated females reject male courtship attempts. In the presence of normal learning and memory, male flies reduce futile courtship behavior in response to sexual rejection by a non-receptive mated female. In this assay, male flies are trained by pairing with a premated female that rejects male courtship attempts. After 1-hour of training, the CI was significantly reduced in the mCherry^RNAi^ control and the less potent Set1^RNAi3^ when compared to naïve flies of the same genotype, indicating normal short-term memory (STM) (Figure 4A, left panel). In contrast, there was no significant reduction in CI after training for Set1^RNAi1^ and Set1^RNAi2^, indicating defective memory upon Set1 knockdown in postmitotic MB neurons. We observed similar results for long-term memory (LTM) (Figure 4A, right panel) with the mCherry^RNAi^ and Set1^RNAi3^ flies showing a significant reduction in CI in response to sexual rejection, and no significant reduction for Set1^RNAi1^ and Set1^RNAi2^. Set1^RNAi1^ and Set1^RNAi2^ showed a significantly reduced LI when compared to the control for both STM and LTM, while Set1^RNAi3^ did not (Figure 4B). Taken together, these data suggest that Set1 is required in postmitotic MB neurons for normal memory.

**Figure 4.**
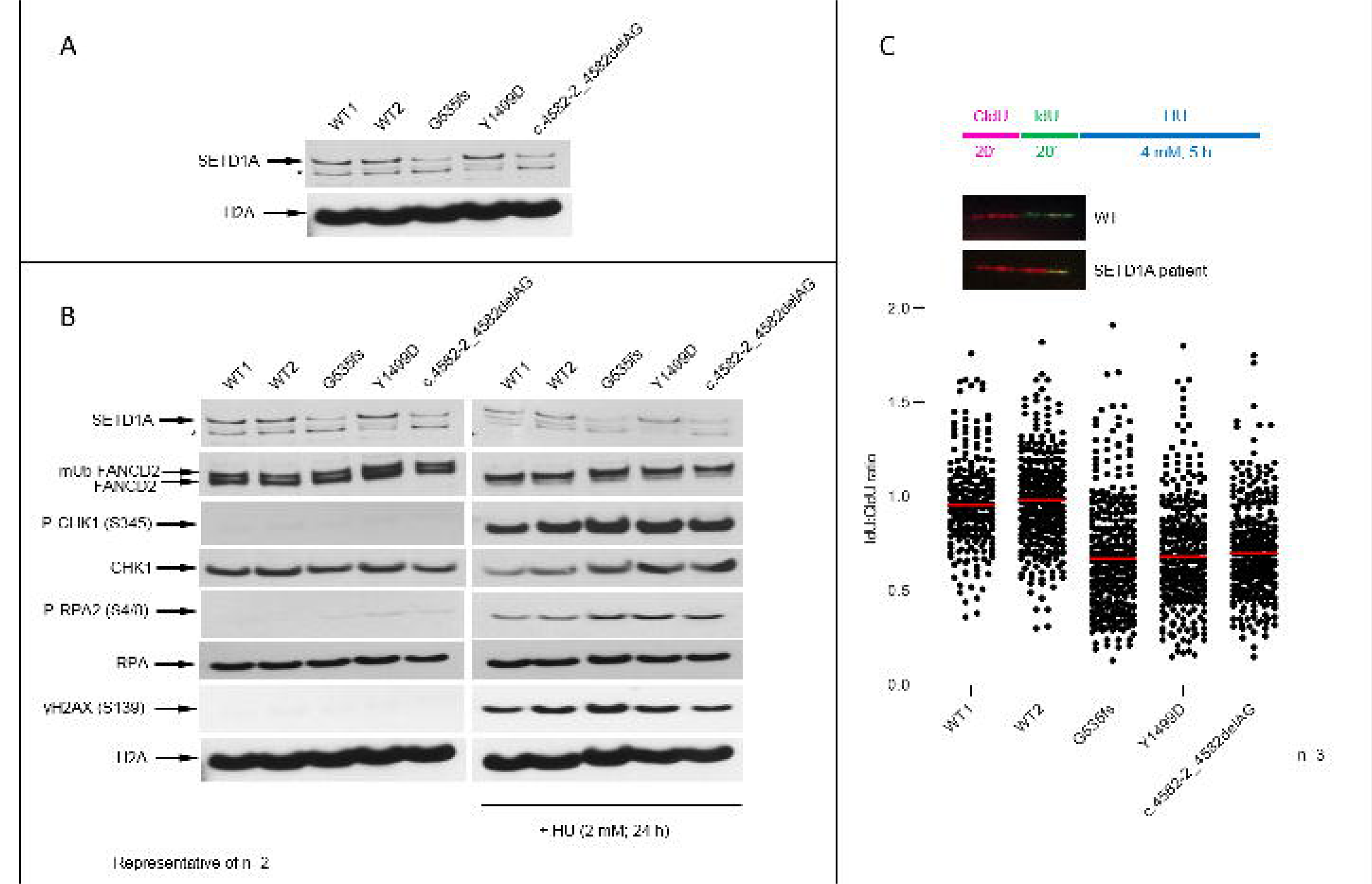
Mushroom body specific knockdown of Set1 causes defects in short- and long-term memory. (A) Courtship memory was measured in control flies and in flies that express Set1 RNAi constructs localized to the mushroom body by the R14H06-Gal4 driver. Three Set1 RNAi lines were compared to the mCherry^RNAi^ control, which is present in the same genetic background as the Set1 RNAi lines. Boxplots show the distribution of Courtship Indices (CIs) for naïve (N) and trained (T) flies of control and Set1 RNAi lines. (*) And (****) indicate a significant reduction in CI trained flies compared to naïve (Mann-Whitney Test; p<0.05 and p<0.0001, respectively). + Indicates the mean. n Is indicated along x-axis. (B) Bar graphs showing the Learning indices (LI), which are single values that are derived from the above CIs according to the formula: LI = (mean CInaïve – mean CItrained)/mean CInaïve). LI was significantly reduced for Set1^RNAi1^ and Set1^RNAi2^, but not Set1^RNAi3^. ** p<0.01, **** p<0.0001, randomization test, 10, 000 bootstrap replicates. Genotypes are indicated below the x-axis.

We also assessed sleep, activity, and baseline courtship behavior in MB-specific Set1 knockdown flies. There was a small but significant (p= 0.03) reduction in sleep for Set1^RNAi1^, however, this reduction was not observed for Set1^RNAi2^ (Supplementary Figure 4A). Overall activity (Supplementary Figure 4B) and naïve courtship levels (Supplementary Figure 4C) were not significantly different in Set1^RNAi1^ and Set1^RNAi2^ when compared to the control. This suggests that courtship conditioning defects are not due to defects in locomotion or baseline courtship ability. It also suggests that memory defects likely arise from the role of Set1 in the MB cells, and not the chordotonal sensory neurons that are also targeted by *R14H06-Gal4*. Disruption of chordotonal organ function would be expected to cause severe defects in balance and locomotion, which clearly are not observed upon *Set1* knockdown with *R14H06-Gal4*.

*R14H06-Gal4* is expressed in the larval MB as well as the adult MB. During larval and pupal development, postmitotic MB neurons undergo morphological changes that are required for normal wiring of the adult fly brain. We asked if memory defects in MB Set1 knockdown flies result from morphological abnormalities arising from defects in MB development. To test this, we labelled R14H06-Gal4 neurons by co-expression of a membrane targeted green fluorescent protein (GFP) and imaged whole mount adult brains using confocal microscopy. We did not observe gross morphological defects in larval, pupal, or adult stages in Set1 knockdown flies (Supplementary Figure 4D). Thus, Set1 appears to be dispensable in postmitotic neurons for normal MB morphogenesis. Taken together with our behavior data, this suggests that Set1 is required for normal functioning of adult MB neurons during the formation of short- and long-term courtship memory.

## Discussion

### Characterization of a Novel Neurodevelopmental Disorder caused by SETD1A Haploinsufficiency

The presented clinical and molecular data strongly support the conclusion that heterozygous LoF of human *SETD1A* causes a novel defined neurodevelopmental disorder. The core features associated with SETD1A haploinsufficiency include global developmental delay or intellectual disability, subtle facial dysmorphisms, and psychiatric problems. In previous studies heterozygous LoF variants in *SETD1A* were observed in individuals with schizophrenia^14^. Therefore, in this study we investigated the behavioral and psychiatric abnormalities that have been observed in our predominantly pediatric cohort. No patient had a previous diagnosis of schizophrenia, but this is expected because of the relatively young age of the patients. With the exception of the very rare childhood onset forms^47^, schizophrenia rarely manifests so early^48^. Interestingly, an adolescent and a young adult from our cohort (age 16 years and 23 years) showed signs that could be linked with schizophrenia or bipolar disorder, though they had no formal psychiatric diagnosis. Combined with the previous reports, this indicates that the germline *SETD1A* variants might predispose individuals to severe neuropsychiatric pathologies later in life. Therefore, this patient group should be carefully monitored for suggestive psychiatric symptoms.

It is notable that in humans all six H3K4 methyltransferases, *SETD1A/B, KMT2A/B*, and *KMT2C/D* have been implicated in neurodevelopmental disorders, though it is also clear that all KMT2 members have distinct molecular functions^30, 41, 49–51^. However, there is an interesting overlap between these chromatin modulating genes and their respective phenotypes (with developmental delay/intellectual disability at its core), based on their shared underlying disease mechanisms^52^.

### *SETD1A* Mutational Mechanisms and Cellular Phenotypes

In the 15 patients investigated in this study we identified 14 different *SETD1A* variants. The majority of these were LoF nonsense or frameshift variants, however a twice recurrent splice-site variant and a missense variant were also identified in the catalytic SET domain, which is required for methyltransferase activity. Previously, SETD1A-mediated H3K4 methylation was shown to be important for protecting against DNA damage at stalled replication forks^12^. We used this cellular phenotype as an output to test the functional effects of clinical *SETD1A* variants in three patient derived lymphoblastoid cell lines. Interestingly, cell lines with heterozygous LoF alleles (p.Gly535Alafs*12, and c.4582-2_4582delAG) showed high levels of nascent DNA degradation, at a level of severity that was similar to that observed in cell lines treated with SETD1A siRNA^12^. Moreover, cells with the heterozygous missense variant (p.Tyr1499Asp) also showed a similar phenotype, thus supporting the pathogenicity of this variant. Taken together, these results suggest that *SETD1A*-associated clinical features likely result from partial loss of SETD1A methyltransferase activity.

In two unrelated patients we found the same *de novo* two base pair (bp) deletion (c.4582-2_4582-1delAG) at the splice acceptor of exon 16. The Human Splicing Finder (version 3.1) predicts that this two bp alteration of the acceptor splice site of exon 16 results in exon skipping^53^. Interestingly, in non-pathogenic situations the overall expression of exon 16 is the highest compared to the other exons of the gene in different tissues^54^ (Supplementary Figure 5). This suggests a significant role for this particular exon for the functionality of the N-SET domain, which is partly encoded by this exon. Our RNA analyses surprisingly revealed that exon skipping did not occur, but the two bp deletion rather resulted in retention of intron 15. In the three other studies describing patients with variants in *SETD1A*, the exact same variant was found in 7 out of the total of 16 patients^14, 23, 24^. In addition, this variant occurs at a frequency of 8.071e^-6^ in the GnomAD database^40^. Authors from the previous studies did not speculate on this seemingly unlikely recurrence of the two bp deletion^14, 24^. However, it raises questions on the frequency and etiology of occurrence of this variant. It is possible that the mutation rate in this genomic area, or specifically around these two base pairs, is relatively high. This may be due to the surrounding sequence homology or the epigenetic state of the DNA, which in turn may allow this deletion to occur. Another explanation might be ascertainment bias, due to the possibility that the specific variant gives rise to ID and/or schizophrenia. Whereas other *SETD1A* variants might also be causative for other phenotypes and therefore would not be included in studies focusing on ID or schizophrenia. In addition, it might be that the c.4582-2_4582-1delAG variant is a so-called selfish mutation. Certain DNA variants that can occur over time in spermatogonial cells can lead to clonal expansion (such as the c.755C>G variant in the *FGFR2* gene in the case of Apert syndrome). This can culminate to the passing of that certain variant from father to child, leading to a higher prevalence in the patient group. Typically these variants are ‘Gain of Function’ (GoF)^55, 56^, so in our cohort this does not seem to be the case.

### The Drosophila SETD1A Orthologue is required in Postmitotic Neurons for Memory

SETD1A and other COMPASS components have been implicated in the regulation of cell proliferation and differentiation. However, there is little evidence that these processes solely underlie the clinical features that we describe in SETD1A haploinsufficiency, as the cognitive deficits and facial dysmorphisms reported in our cohort are mild. Defects in proliferation or differentiation of neural progenitors might have more dramatic effects on structure and function of the brain, and cause profound cognitive deficits or lethality. In line with this, we identified a role for *Drosophila* Set1 in the regulation of memory formation in postmitotic neurons. This finding is consistent with studies demonstrating a role for different histone modifications in the epigenetic regulation of memory^57^. For example, H3K4me3 is dynamically regulated in the rodent hippocampus during memory formation and Setd1a^+/-^ mice exhibit deficits in working memory^58, 59^. Several other H3K4 methyltransferases have also been implicated in memory, including mouse Kmt2a/Kmt2b and *Drosophila* trr^30, 49, 60^. Thus, our findings on *SETD1A* and *Drosophila* Set1 support a growing body of literature highlighting the conserved role of H3K4 methylation in regulating the function of adult neurons post development.

The specific role of H3K4 methylation in memory formation is not known. Set1-related proteins might regulate several different aspects of postmitotic neuronal biology that could all impact memory. For example, we have shown here that patient cell lines have elevated damage in nascent DNA. Terminally differentiated neurons are often exposed to DNA damage and an efficient response is critical, as these neurons are not easily replaced. Additionally, the Set1 protein family might be important for maintaining the cell-type specific gene expression programs that would be critical for normal functioning of memory neurons. Set1-related proteins might also be required for induction of learning-induced gene expression changes that are known to be required for long-term memory formation.

Despite the lack of a precise pathogenetic mechanism, the finding that Set1 regulates memory post development could have implications for therapies. Further understanding of SETD1A function in the postnatal brain could open up possibilities for therapy that would not be possible if pathogenic mechanisms occurred entirely during the prenatal stages of development.

### Compliance with ethical standards

All procedures performed in studies involving human participants were in accordance with the ethical standards of the institutional and/or national research committee and with the 1964 Helsinki declaration and its later amendments or comparable ethical standards.

Informed consent was obtained from all individual participants included in the study as part of the diagnostic workflow. Additional informed consent was obtained from all individual participants for whom identifying information is included in this article.

## Supporting information

Supplementary Figure 5 SETD1A expression in tissue

Supplementary Table 1 Clinical Patient Overview

Supplementary Figure 1 Metadome Tolerance Landscape

Supplementary Figure 2 Intron Retention By AG Deletion

Supplementary Figure 3 Age Distribution Patient Cohort

Supplementary Figure 4 Drosophila Results

Supplementary Data 2 Results

## Acknowledgements

We are grateful to all individuals and their parents for participating in this study.

This work was supported by the Netherlands Organization for Health Research and Development (ZonMw grant 91718310) to T.K., ERA-NET NEURON-102 SYNSCHIZ (grant 013-17-003 4538) to D.S., Canadian Institutes of Health Research, Canadian Foundation for Innovation, and the Canada Research Chairs Program to J.M.K., MRC Career Development Fellowship (MR/P009085/1) and a University of Birmingham Fellowship to M.R.H.

We thank the Bloomington Drosophila Stock Center at Indiana University for providing all *Drosophila* strains used in this study.

Sequencing and analysis for patient 2 were provided by the Broad Institute of MIT and Harvard Center for Mendelian Genomics (Broad CMG) and was funded by the National Human Genome Research Institute, the National Eye Institute, and the National Heart, Lung and Blood Institute grant UM1 HG008900 and in part by National Human Genome Research Institute grant R01 HG009141.

## Conflict of interest

All contributors have read and approved the submission to the journal.

M.T. Cho and K. McWalter are employees of GeneDx. We are not aware of any other conflict of interest.

Supplementary information is available at MP’s website.

## Supplementary information

Supplementary Data 1 – Methods

Detailed description of used methods with cell culture, immunoblotting, antibodies, RNA isolation and acquisition of fly stocks, and courtship conditioning.

Microsoft Word file

Supplementary Data 2 – Results

Additional clinical information, and additional highlighted individual cases.

Microsoft Word file

Supplementary Table 1

Clinical features of 15 novel described patients with *SETD1A* pathogenic variants and a summary of 16 previously described patients with *SETD1A* variants by Singh *et al.*^14^.

Abbreviations: NA: Not available; NR: Not Reported; SD: Standard Deviation. 1. SHPRH: unknown variant, *de novo*; 2. 16p13.11dup: non-TRIO sequenced; 3. POLG: c.2740A>C, *de novo*; 4. CNTNAP2: unknown variant, maternally inherited; 5. EFNB2: c.876_884dup p.(Leu293_Thr295dup), *de novo*; 6. NHS: c.3316C>T p.(His1106Tyr), mother carrier.

Text summary

Detailed description of the clinical features and molecular diagnosis of patients with *SETD1A* pathogenic variants and a summary of 16 previously described patients with *SETD1A* variants by Singh *et al.*^14^.

Microsoft Excel file

Supplementary Figure 1

Metadome tolerance landscape. By using homologous domain relations and combining data from GnomAD and ClinVar this online data platform produces a ‘tolerance landscape’ of a gene, from which a degree of tolerance of certain variants can be determined.

Supplementary Figure 2

*SETD1A* c.4582-2_4582-1del effect on splicing causing COMPASS complex Set1 subunit interruption and mRNA degradation.

Supplementary Figure 3

Bar graph showing the age distribution of patient cohort. The absolute age of the patients in years at the time of inclusion. The range of age distribution is 2 – 23 years, with a mean of 5 years.

Supplementary Figure 4

Sleep, activity, courtship behavior, and development are not generally perturbed in MB-specific Set1 knockdown flies. Graphs showing sleep (A), activity (B), and naïve courtship indices (C) in flies expressing Set1 or mCherry UAS-RNAi constructs with R14H06-Gal4. Generally these behavioral parameters are unaffected by Set1 knockdown, however, Set1^RNAi1^ flies did show a small but significant reduction of sleep. Error bars indicate SEM. Sleep and activity were compared using AONVA. CIs were compared by Kruskal Wallis. P-values < 0.05 are indicated. (D) Confocal projections of adult fly mushroom bodies expressing GFP show morphological similarity between mCherry^RNAi^ (n=11), Set1^RNAi1^ (n=16), and, Set1^RNAi2^ (n=14).

Supplementary Figure 5

Exon expression of *SETD1A* in tissue according to GTEx Portal. The median read count per base indicates that the expression of exon 16 in tissue is overall the highest. This implicates an important role of this particular exon in the functionality of the gene.

