## Supplementary figures and images for "Characterization of *SETD1A* haploinsufficiency in humans and *Drosophila* defines a novel neurodevelopmental syndrome"

### Supplementary Figure 1 Metadome Tolerance Landscape

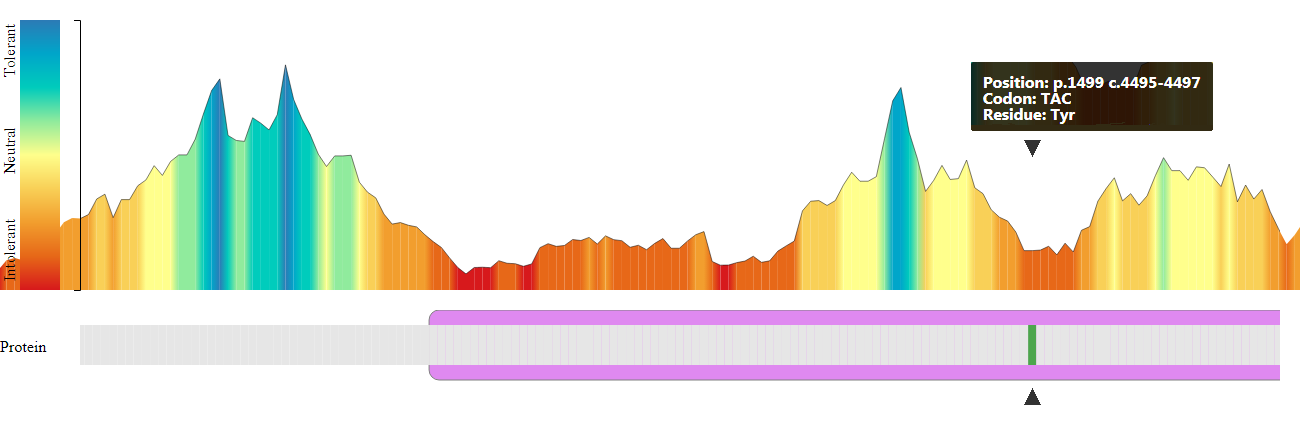

### Supplementary Figure 2 Intron Retention By AG Deletion

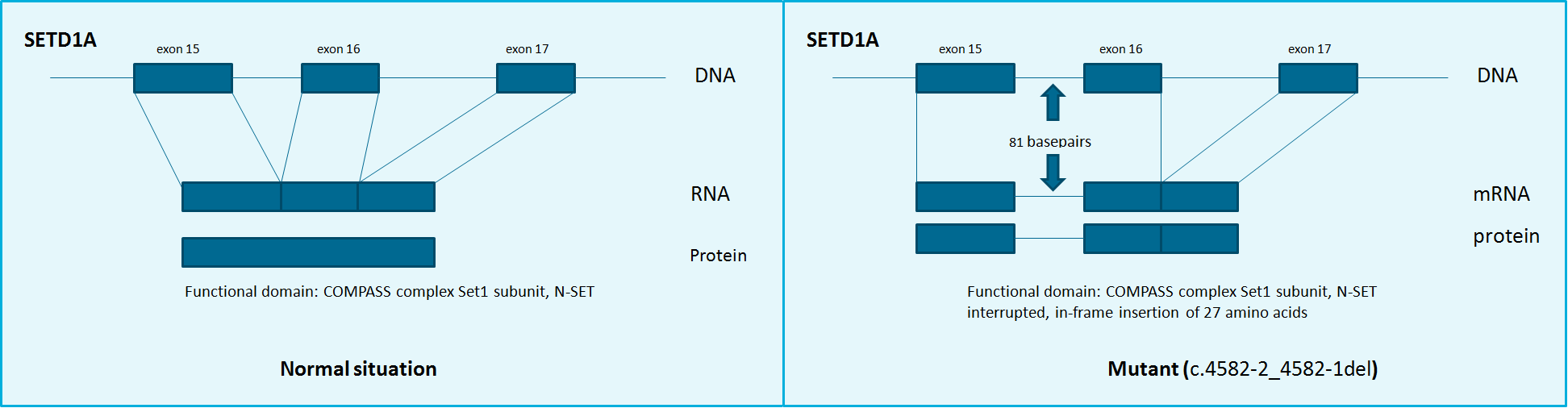

### Supplementary Figure 3 Age Distribution Patient Cohort

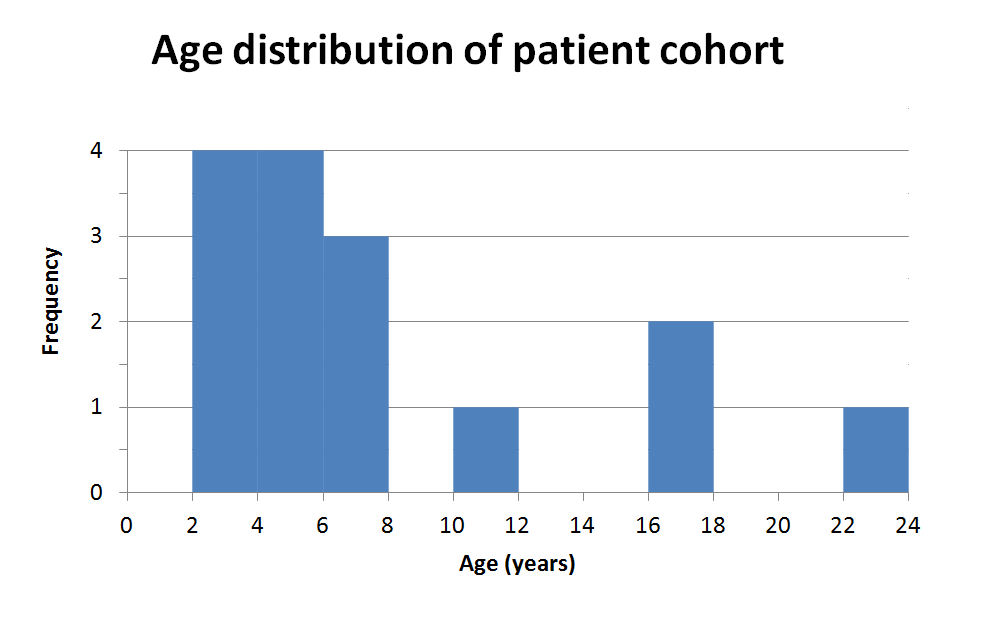

### Supplementary Figure 4 Drosophila Results

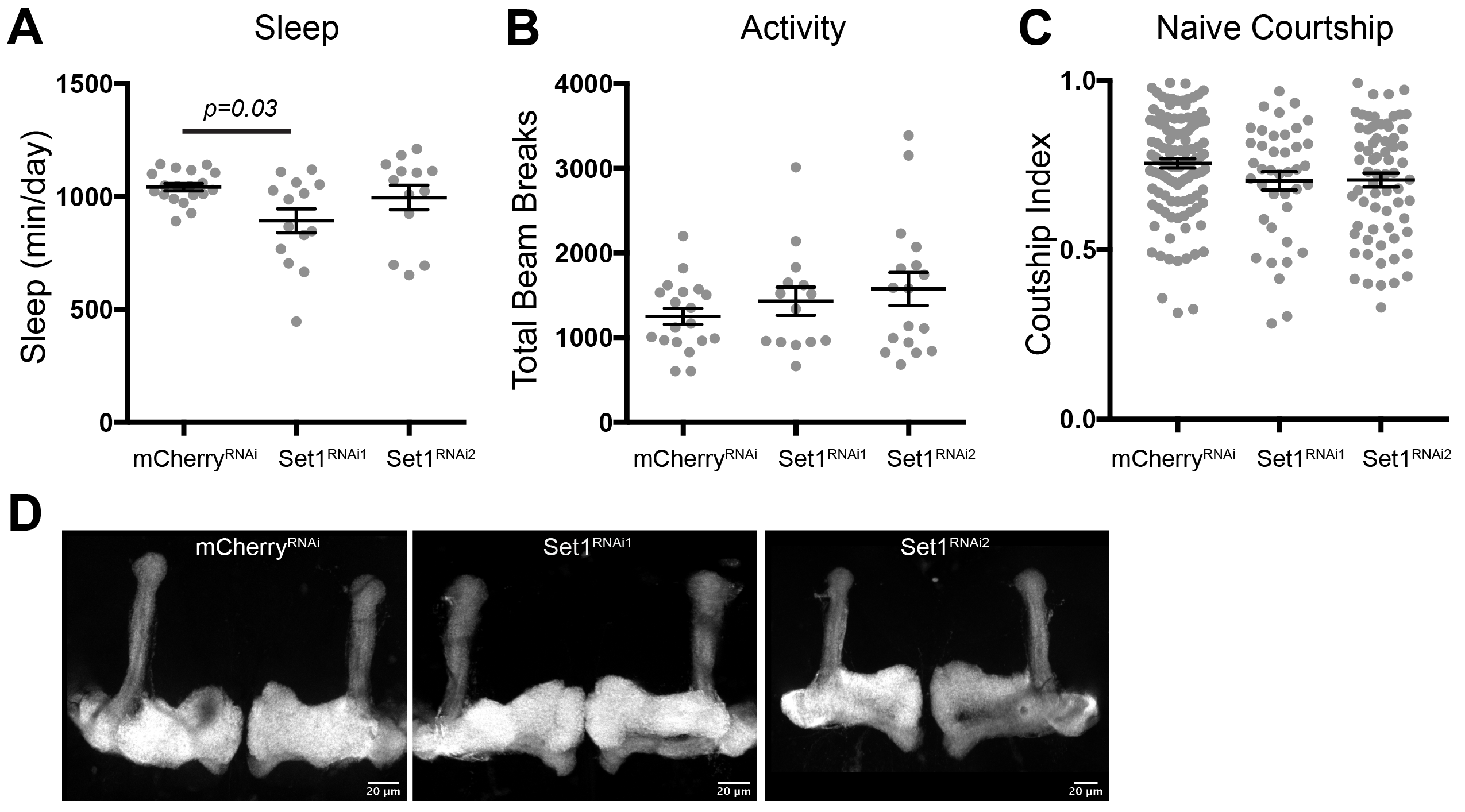

### Supplementary Figure 5 SETD1A expression in tissue

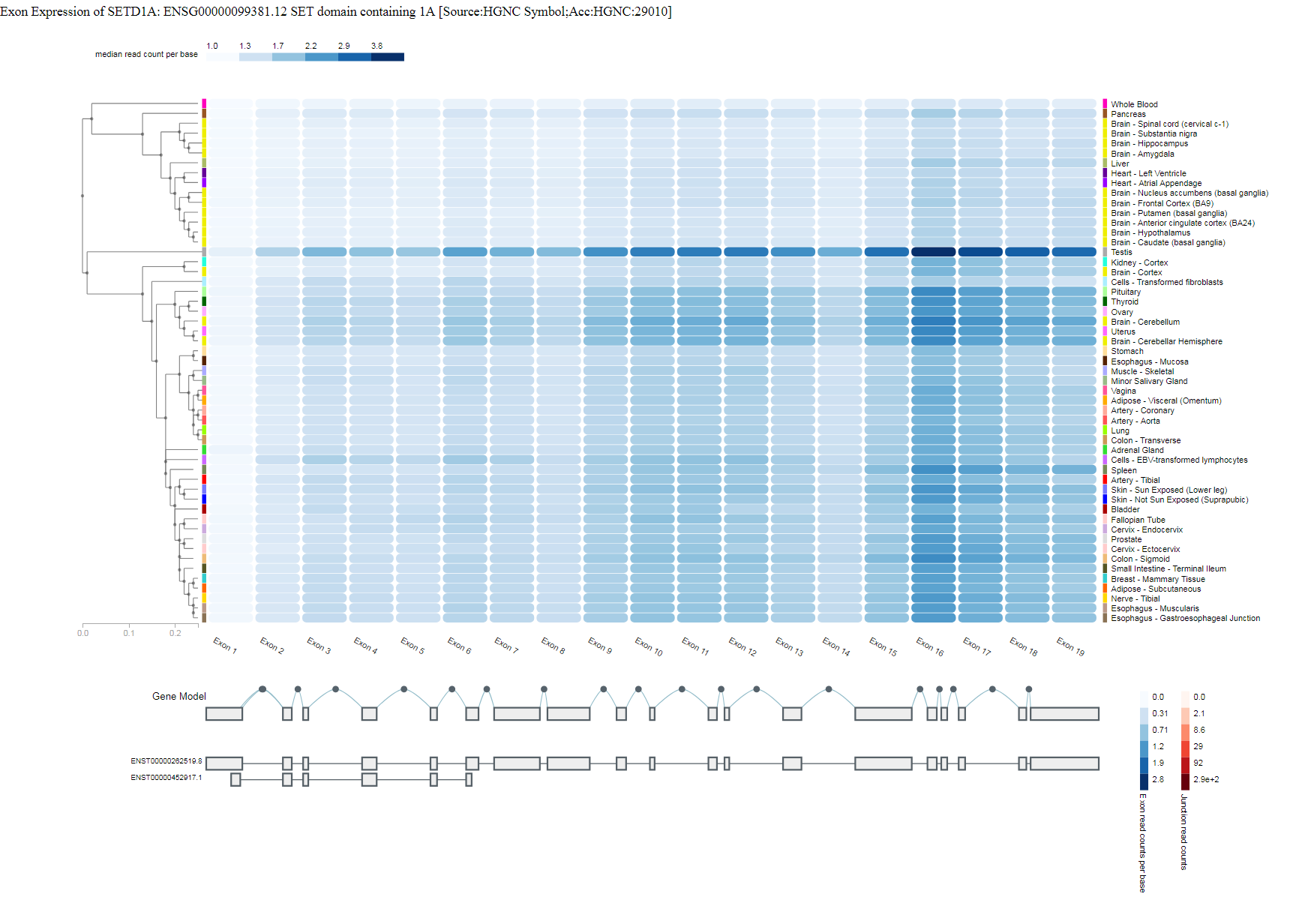
