## Supplementary Table 1 Clinical Patient Overview for "Characterization of *SETD1A* haploinsufficiency in humans and *Drosophila* defines a novel neurodevelopmental syndrome"

| Reference<br>Patient | Novel *SETD1A* variant cases |  |  |  |
| --- | --- | --- | --- | --- |
|  | 1 | 2 | 3 | 4 |
| Gender | Male | Female | Female | Male |
| Age of examination | 2 Years 10 Months | 6 Years | 16 Years 6 Months | 6 Years 8 Months |
| Molecular diagnosis |  |  |  |  |
| Genomic position (NG_052948.1) (Hg19: Chr16) | g.30970161C>T | g.30976077dup | g.30976205_30976208del | g.30976426G>T |
| cDNA change (NM_014712.2) | c.109C>T | c.1014dupC | c.1144_1147del | c.1363G>T |
| Protein change (NP_NP_055527.1) | p.Gln37* | p.Ala339Argfs*23 | p.(Tyr382Hisfs*114) | p.Glu455* |
| Mutation type | Nonsense | Frameshift | Frameshift | Nonsense |
| Inheritance | De novo | De novo | De novo | De novo |
| Additional variant(s) | - | - | - | - |
| Pregnancy/delivery |  |  |  |  |
| Pregnancy | Mother took gabapentin and birth control first 12 weeks | History of miscarriage | History of miscarriage | Uncomplicated |
| Delivery | Caesarean section | Uncomplicated | Caesarean section | Uncomplicated |
| Gestational age | Full term | Full term | Full term | Full term |
| Birth weight (g) | 3033 (-1.3 SD) | NR | 2722 (-0.4 SD) | 3320 (+0.1 SD) |
| Growth |  |  |  |  |
| Height (cm) | 91.4 (-1.4 SD) | 126 (+2 SD) | 159.5 (-1.4 SD) | 106 (-3.4 SD) |
| Weight (kg) | 12.6 (-1.6 SD) | 27.4 (+1.5 SD) | 69.8 (+1.5 SD) | 19 (-1.7 SD) |
| BMI (kg/m<sup>2</sup>) | 15.1 (-0.9 SD) | 17.3 (+1.1 SD) | 27.4 (+1.4 SD) | 16.9 (+0.9 SD) |
| Head circumference (cm) | 48.3 (-1.2 SD) | NA | 56 (+0.5 SD) | 53.2 (at 3 Years 4 Months) (+1.4 SD) |
| Development |  |  |  |  |
| Intellectual disability/Learning difficulties | + | - | + | + |
| Global developmental delay | + | - | + | + |
|  | +, Does not follow |  |  | +, More severely |
| Delayed speech and language commands |  |  | + | affected than motor |

|  |  |  |  |  |  |
| --- | --- | --- | --- | --- | --- |
|  |  | +, Delayed gross and fine motor development |  |  |  |
|  | Motor delay |  |  | + | + |
| Behavioral/ psychiatric abnormalities |  |  |  |  |  |
|  | Behavioral abnormality | + | + | + | - |
|  | Sleep disturbance | NA | - | - |  |
|  | Other | Autistic behaviour;<br>Problems with<br>transitions, problems<br>with food textures | - | Attention deficit<br>hyperactivity disorder;<br>Short attention span;<br>Hears voices telling to<br>do things and seeing<br>things |  |
| Neurologic abnormalities |  |  |  |  |  |
| Seizures |  | - | +, Onset at 4.5 years.<br>Epilepsy syndrome<br>consistent with Doose<br>Syndrome. | +, Benign focal onset<br>seizures in early<br>childhood | - |

|  |  |  |  |  |
| --- | --- | --- | --- | --- |
| Hypotonia | + | - | - | + |
| Morphological CNS abnormalities | - | - | - | +, Chiari I malformation |
| Other | - | Encephalopathy;<br>Dysarthria (mild);<br>Tremor (slight);<br>Dysgraphia (right hand) | Deficit in phonologic short-term memory due to anxiety?; EEG abnormality | - |
| Musculoskeletal abnormalities |  |  |  |  |
| Joint hypermobility | - | - | - | + |
| Abnormality of the extremities | +, Slightly broad/wide fingers | - | - | +, Full, puffy hands and feet; Mild 2-3 syndactyly feet; Broad halluces |

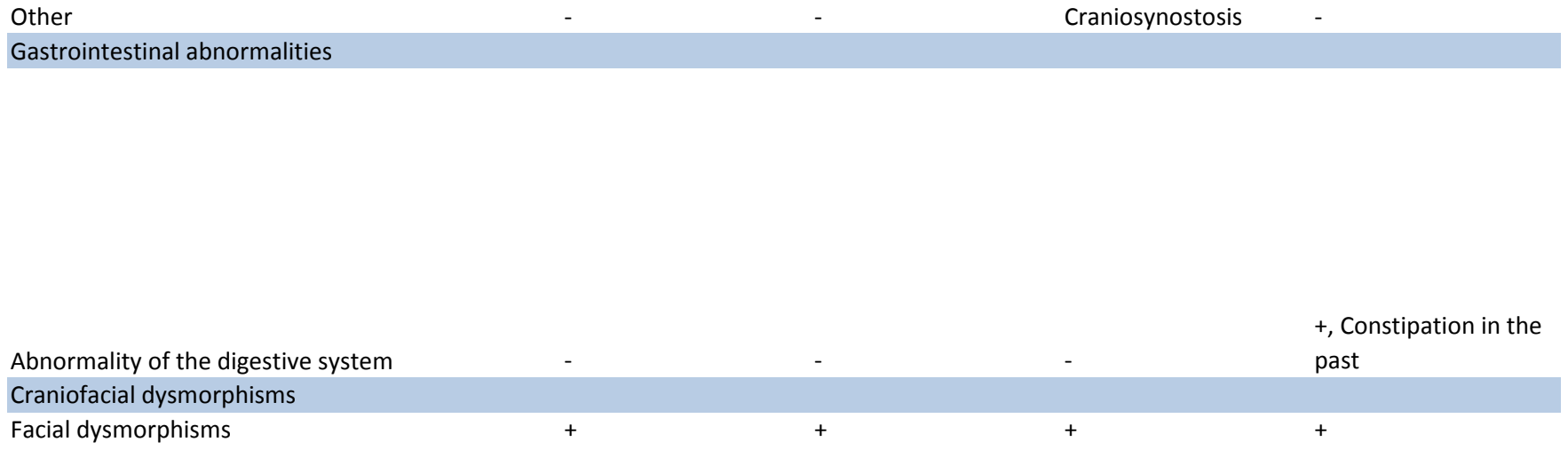

Epicanthus; Narrow palpebral fissures; Slightly upturned nose; Thickened alae nasae and columella; Thick vermilion borders; Wide mouth; Widely spaced teeth

Epicanthus; Hypertelorism; Wide nose; Wide mouth

Tall protruding forehead; Receded anterior hairline; Downslanted palpebral fissures; Cupped ear; Low-set ears; Prominent chin

Mild hypertelorism; Low nasal bridge; Downslanted palpebral features; Peri-orbital fullness; Mild ptosis; Full lips (These dysmorphisms were such that Coffin-Lowry syndrome was specifically looked for); Cupped ears; Minimal notched alae nasi (The dysmorphisms around the eyes and the full lips have greatly improved over time. As has the puffiness of hands and feet)

Other clinical features

Recurrent otitis
media; Recurrent
pneumonia; Myopia;
Astigmatism; Easy
fatigability

Recurrent middle ear
infections;
Abnormality of
refraction

| 5 | 6 | 7 | 8 | 9 | 10 | 11 |
| --- | --- | --- | --- | --- | --- | --- |
| Male<br>4 Years 4 Months<br>Deceased | Female<br>10 Years 4 months | Female<br>3 Years 6 Months | Female<br>3 Years 9 Months | Female<br>5 Years 2 months | Female<br>17 Years 1 Month | Male<br>3 Years |
| g.30976558C>T | g.30976665_30976666del | g.30966169 | g.30978864G>T | g.30980962C>T | g.30991044_30991054del | g.30991089_30991090del |
| c.1495C>T | c.1602_1603del | c.2288_2289insA | c.2725G>T | c.2968C>T | c.3937_3947del11 | c.3982_3983del |
| p.Gln499* | p.Gly535Alafs*12 | p.Val764SerfsTer*61 | p.Glu909* | p.Arg990* | p.Pro1313Argfs*17 | p.Phe1328Glnfs*5 |
| Nonsense | Frameshift | Frameshift | Nonsense | Nonsense | Frameshift | Frameshift |
| De novo | De novo | De novo | De novo | De novo | Unknown (not maternal) | De novo |
| SHPRH (1) | - | - | 16p13.11dup (2);<br>RBBP8 c.1931del | - | - | POLG (3) |
| Uncomplicated | Uncomplicated;<br>History of<br>miscarriages | Uncomplicated | Uncomplicated | Uncomplicated<br>Caesarean section due<br>to breech<br>presentation | Oligohydramnios (at<br>39 weeks); Maternal<br>obesity | Uncomplicated |
| Uncomplicated | Fetal emergency<br>Caesarean section | Caesarean section | Uncomplicated |  | Uncomplicated | Uncomplicated |
| Full term | Full term | Full term | Full term | Full term | Full term | Premature (34 weeks) |
| 2410 (-2.9 SD) | Normal | 3200 (-0.6 SD) | 3050 (-0.7 SD) | 3240 (-0.6 SD) | 2948 (-1.2 SD) | 2230 (0 SD) |
| Normal |  |  |  |  |  |  |
| 85 (<-2.5 SD) | 146 (0 SD) | 102 (+0.2 SD) | 106 (+0.7 SD) | 110 (- 1 SD) | 158.8 (-1.6 SD) | 98 (0 SD) |
| 14 (-2.3 SD) | 44.6 (+ 1.6 SD) | 16 (+0.1 SD) | 22.7 (+2.6 SD) | 20 (+ 1 SD) | 100.5 (+4.2 SD) | 18 (+ 1.5 SD) |
| 19.4 (+1.9 SD) | 20.9 (+1.2 SD) | 15.4 (-0.1 SD) | 20.2 (+2.4 SD) | 16.5 (+0.9 SD) | 39.9 (+2.3 SD) | 18.7 (+2.0 SD) |
| 49 (-1.4 SD) | 55.8 (+ 1.7 SD) | 54 (+2.7 SD) | 49.5 (-0.3 SD) | 53.5 (+ 2 SD) | 55.8 (+0.4 SD) | NA |
| + | + | + | - | + | + | + |
| + | + | + | + | + | + | + |
|  |  | +, Approximately 6<br>months behind | + | + | + | +, Limited vocabulary<br>and problems with<br>pronunciation |
| + | +, Rigid movements | +, Approximately 6 months behind | + | + | - | + |
| + | + | + | + | + | + | + |
| NA | - | +, Wakes up frequently | + | - | +, Requires very little sleep | + |
| Sensory aversions | Anxiety; Short attention span; Overfriendliness (affectionate behaviour); Eating difficulties in line with Anorexia | - | Overfriendliness (very open and social); Phonophobia | Anxiety; Overfriendliness | Autism Spectrum Disorder; Aggressive behaviour; Skin-picking; Pica | Aggressive behavior; Short attention span; Stubborn |

|  |  |  |  |  |  |  |
| --- | --- | --- | --- | --- | --- | --- |
| - | - | - | - | - | - | + , Severe since 2 years age |

|  |  |  |  |  |  |  |
| --- | --- | --- | --- | --- | --- | --- |
| + | + | - | - | + | - | + |
| +, T2 Hyperintensity | NA | - | NA | - | +, Abnormalities of the cerebral white matter | NA |

Generalized EEG slowing, no epileptiform activity

|  |  |  |  |  |  |  |
| --- | --- | --- | --- | --- | --- | --- |
| - | - | - | - | - | - | - |
| - | + | +, Extremely hypermobile | + | + | + | NA |

|  |  |  |  |  |  |  |
| --- | --- | --- | --- | --- | --- | --- |
| +, Tapered fingers | +, Long fingers | +, Tapered fingers | +, Pes planus (mild) | +, Pes planus | +, Pes planus; Achilles tendon contractures | +, Pes planus |
| - | Hip dysplasia | - | Kyphosis; Neonatal<br>plagiocephaly<br>(treated) | - | - | Hemiplegia left, due<br>to epilepsy treatment |

|  |  |  |  |  |  |  |
| --- | --- | --- | --- | --- | --- | --- |
| +, Gastrojejunal tube<br>feeding; Constipation;<br>Failure to thrive | - | +, Insatiable appetite | +, Gastroesophageal<br>reflux (in neonatal<br>period) | +, Feeding slowly;<br>Constipation | +, Frequent bowel<br>movements | NA |
| + | + | + | + | + | + | + |

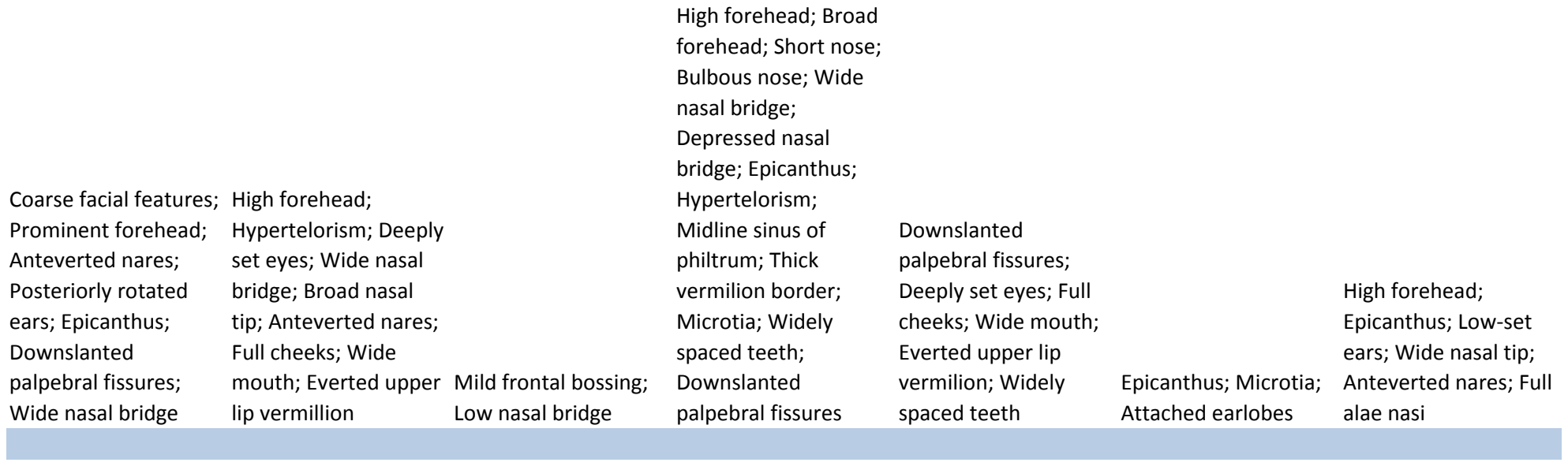

Common variable
immune deficiency;
Decreased antibody
level in blood;
Pulmonary
hypertension; Chronic
interstitial pulmonary
disease;
Tracheostomy/
Ventilator dependent

Sensitive skin

Refractional
abnormality;
Strabismus;
Respiratory problems
at birth

Recurrent infections
(ear and respiratory);
Bruising susceptibility;
Strabismus (small)

Visual impairment;
Hemangioma (left
knee)

Leukocytosis
(suspected to be
benign); Acne
(pimples/comedones)
at back and buttocks;
Acanthosis nigricans;
Pre-diabetics

-

|  |  |  |  |  | Previously reported<br>*SETD1A* variant cases<br>by Singh *et al.* (6) |
| --- | --- | --- | --- | --- | --- |
| 12 | 13 | 14 | 15 | Total novel *SETD1A*<br>variant cases<br>1-15 (15 patients) | 16-32 (16 patients) |
| Male | Male | Female | Male | Female (8/15); Male<br>(7/15) | NR |
| 7 Years | 4 Years 7 Months | 5 Years 10 Months | 23 Years | 2 Years 10 Months -<br>23 Years | NR |
| g.30991804A>G | g.30991892T>G | g.30992058_3099205<br>9del | g.30992058_3099205<br>9del | c.4582-2_4582delAG<br>(2/15) | c.4582-2_4582delAG<br>(7/16) |
| c.4409-2A>G | c.4495T>G | c.4582-2_4582delAG | c.4582-2_4582delAG |  |  |
| p.(?) | p.Tyr1499Asp | p.(?) | p.(?) |  |  |
| Splice acceptor | Missense | Splice acceptor | Splice acceptor | Frameshift (6/15);<br>Nonsense (5/15);<br>Missense (1/15);<br>Splice acceptor (3/15) | LoF (16/16) |
| De novo | De novo | De novo | De novo | De novo (14/14) | De novo (6/16); Case<br>(8/16); Maternally<br>inherited (2/16) |
| CNTNAP2 (4) | EFNB2 (5); NHS (6) | - | - | Additional variants<br>(5/15) | NR |
| Uncomplicated | Growth retardation;<br>Dizygotic twin pregnancy | Intra uterine growth retardation;<br>lkaclomide induced pregnancy | Unknown | History of miscarriage (3/14); Growth retardation (2/14); Twin-pregnancy (1/14); Oligohydramnios (1/14); Maternal obesity (1/14) |  |
| Caesarean section | Breech presentation | Uncomplicated | Unknown | Caesarean section (6/14)<br>Premature birth (1/14) | Breech presentation (1/16) |
| Full term | Full term | Full term | Unknown |  |  |
| 3200 (-0.6 SD) | 2020 (< -1 SD) | 2470 (-2.7 SD) | (> 2 SD) | Large for gestational age (1/15); Small for gestational age (1/15) |  |
| 128 (+1.3 SD) | 103.5 (-1.5 SD) | 106 (-2.2 SD) | 187 (+0.6 SD) | Tall stature (1/15);<br>Short stature (3/15) | Overgrowth (1/16);<br>Short stature (1/16) |
| 27.8 (+1.3 SD)<br>17 (+0.8 SD) | 15.7 (-1.6 SD)<br>14.7 (-0.8 SD) | 17.5 (-1.4 SD)<br>15.6 (+0.3 SD) | 102.8 (>+2.5 SD)<br>29.4 (+1.5 SD) | Overweight (3/15) | Truncal obesity (1/16) |
| 52.5 (+0.4 SD) | 49.5 (-1.1 SD) | 50.7 (-0.1 SD) | 62 (+2.5 SD) | Macrocephaly (3/13) | Macrocephaly (1/16) |
| + | + | - | + | + (12/15) | + (6/7) |
| + | + | + | + | + (14/15) | + (7/16) |
| + | + | +, Speech apraxia | + | + (14/14) | + (6/7) |

+, Delayed gross and
fine motor
development;

Clumsiness

+

+

+

+ (13/14)

+ (6/7)

+

+

+

+

+ (14/15)

+ (13/16)

+, Night terrors;

Insomnia

+

-

+, As infant

+ (7/12)

+, Night terrors (1/13)

|  |  |  |  |  |  |  |  |
| --- | --- | --- | --- | --- | --- | --- | --- |
| Anxiety; Aggressive behaviour; Short attention span; Temper tantrums; Irritability; Obsessive-compulsive traits |  | Anxiety; Aggressive behaviour; Temper tantrums |  | Impaired social interactions | Aggressive behaviour; Addictive behaviour; PDD-NOS; Bipolar disorder like symptoms; Short attention span | Aggressive behaviour (5/14); Short attention span (5/14); Anxiety (4/14); Autistic behaviour (3/14); Overfriendliness (3/14); Temper tantrums (2/14); Phonophobia (1/14); Sensory aversions (1/14); Obsessive-compulsive traits (1/14); Addictive behaviour (1/14); Attention deficit hyperactivity disorder (1/14); Bipolar disorder like symptoms (1/14); Anorexia like behaviour (1/14); Self-injurious behaviour (1/14); Pica (1/14) | Schizophrenia (10/13); Psychotic episodes (5/13); Impaired social interaction (3/13); Obsessive-compulsive behavior (2/13); Aggressive behavior (2/13); Mental deterioration (2/13); Anxiety (1/13); Diminished motivation (1/13) |

|  |  |  |  |  |  |
| --- | --- | --- | --- | --- | --- |
| - | - | - | - | + (3/15) | + (2/16), Grand mal status epilepticus (1/2) |

|  |  |  |  |  |  |
| --- | --- | --- | --- | --- | --- |
| + | + | + | - | + (9/15) | + , Infantile axial hypotonia (1/16) |
| - | - | + , Abnormality of the corpus callosum | - | + (4/12) |  |
|  |  |  |  | EEG abnormality (2/15); Encephalopathy (1/15); Dysarthria (1/15); Tremor (1/15); Dysgraphia (1/15); Abnormal downward eye movements (1/15); Sensory processing disorder (1/15); Phonological short-term memory deficit (1/15) |  |
| - | - | Abnormal downward eye movements; Sensory processing disorder | - |  | NR |
| - | - | + | NA | + (7/13) | NR |
|  |  |  |  | + (12/15); Pes planus (6/12); Broad/wide fingers (3/12); Long fingers (2/12); Tapered fingers (2/12); Achilles tendon contractures (1/12); Mild 2-3 syndactyly feet (1/12) |  |
| + , Long fingers; Prominent fingertip pads; Joint contracture of the 5th finger bilateral; Sandal gap |  |  |  |  |  |
|  | + , Pes planus (mild) | + , Pes planus | - |  | NR |
| Pectus excavatum of inferior sternum (mild) | Craniosynostosis; Neonatal plagiocephaly; Hyperlordosis | Postural instability; Broad-based gait; Right-sided preference; Hyperlordosis; Congenital torticollis | - | Plagiocephaly (2/15); Hyperlordosis (2/15); Craniosynostosis (2/15); Kyphosis (1/15); Pectus excavatum (1/15); Hip dysplasia (1/15); Postural instability (1/15); Right sided preference (1/15); Congenital torticollis (1/15) | NR |
| NA | + | +, Feeding difficulties during breast feeding | NA | + (8/12); Feeding difficulties (3/8); Constipation (3/8); Gastroesophageal reflux (1/8); Frequent bowel movements (1/8); Insatiable appetite (1/8) | NR |
| + | + | + | + | + (15/15) | + (5/16) |

|  |  |  |  |  |
| --- | --- | --- | --- | --- |
| Narrow forehead; |  |  |  | High forehead (7/15); |
| Deeply set eyes; |  |  |  | Low-set ears (4/15); |
| Hypertelorism, |  |  |  | Microtia (3/15); |
| Downslanted |  |  |  | Downslanted |
| palpebral fissures; |  |  |  | palpebral fissures |
| Wide nasal bridge and |  |  |  | (6/15); Epicanthus |
| base; Broad nasal tip; |  |  |  | (7/15); Deeply set |
| Anteverted nares; |  |  |  | eyes (4/15); |
| Broad philtrum; |  |  |  | Hypertelorism (6/15); |
| Narrow mouth; Thick | Pominent suture right; |  |  | Wide nose (6/15); |
| upper and lower lip | High forehead; Broad |  |  | Anteverted nares |
| vermillion; Tented | forehead; Mild |  |  | (4/15); Full cheeks |
| upper lip vermillion; | hypertelorism; Tented |  |  | (2/15); |
| Protruding and low | upper lip vermillion; |  | High forehead; Tented | Everted/tented upper |
| set ears; | Ankyloglossia; |  | upper lip vermillion; | lip vermillion (5/15); |
| Underdevelopment of | Abnormality of the |  | Deeply set eyes; | Wide mouth (4/15); |
| the helix | dentition | Epicanthus | Microtia; Low set ears | Widely spaced teeth |
|  |  |  | (3/15) |  |

|  |  |  |  |  |  |
| --- | --- | --- | --- | --- | --- |
| Low-frequency sensorineural hearing impairment | Soft skin; Recurrent infections (ear) | Myopia; Recurrent acute otitis media; Hearing impairment due to tympanic membrane perforation | Asthma; Recurrent ear infections | Recurrent infections (7/15); Skin abnormalities (4/15); Hemangioma (1/15); Visual impairment (6/15); Hearing impairment (2/15); Leukocytosis (1/15); Pre-diabetics (1/15); Easy fatigability (1/15); Respiratory problems at birth (1/15) | Capillary hemangiomas (2/16); Wide intermammary distance (1/16); Breath holding spells (1/16); Renal duplication (1/16); Hydrocele testis (1/16) |

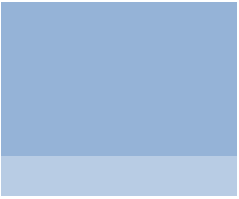

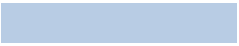

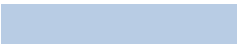

100%

100%

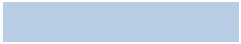

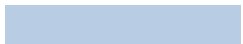

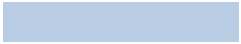

\_\_\_\_\_

\_\_\_\_\_

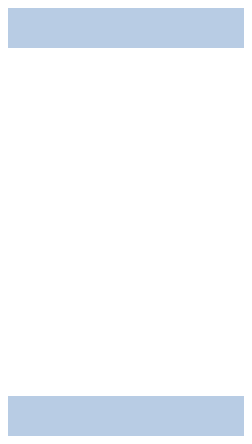

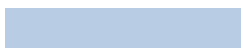
