## Supplementary Data 1 Methods for "Characterization of *SETD1A* haploinsufficiency in humans and *Drosophila* defines a novel neurodevelopmental syndrome"

### Cell Culture

Lymphoblastoid cell lines (LCLs) from 3 individuals with *de novo* SETD1A variants or 2 unaffected counterparts were created by transforming peripheral blood lymphocytes with EBV. Cells were cultured in RPMI-1640 with 15% fetal bovine serum (Life Technologies).

### Immunoblotting

For western blotting, whole cell extracts of LCL were obtained by lysis in UTB buffer (8 M Urea, 50 mM Tris, 150 mM β-mercaptoethanol, protease inhibitor cocktail (Roche)). Cell extracts were sonicated, clarified by centrifugation, and protein concentration was determined by Bradford Assay (Bio-Rad). Polypeptides were separated by SDS-PAGE, transferred onto nitrocellulose membrane, incubated with primary antibody overnight, followed by HRP-linked secondary antibody for 1hr at room temperature. The signal was detected using ECL western blotting substrate (GE Healthcare).

### Antibodies

The antibodies used were: RPA2 (Calbiochem); Chk1, FANCD2, (Santa Cruz Biotechnology); SETD1A, phospho-RPA2 Ser-4/8, phospho-Chk1 Ser-345 (Bethyl); H2A, γ-H2AX (Merck); BrdU (CldU) (Bio-Rad); BrdU (IdU) (Becton Dickinson); anti-mouse-HRP, anti-rabbit-HRP (Dako), Alexa-Fluor anti-rat 555 Alexa-Fluor anti-mouse 488 (Thermo Fisher).

### RNA Isolation

To study the effect of the splice site mutation in patient 15, RNA was isolated from EBV derived cell lines, with and without Cycloheximide from EBV cell lines from this patient using the QIAamp RNA Blood Mini kit from Qiagen. cDNA synthesis and RT-PCR were carried out using the Titan One Tube RT-PCR kit from Sigma-Aldrich. Target specific primers were used to amplify the target locus exon 15-16 (*SETD1A*, NM_014712.2).

Forward primer ACGCAGTGAGTTTGAACAGA (exon 14),

Reverse primer GTGCAGCAGTGGTTGATGAA (exon 18).

The amplified PCR-products were purified and sequencing was performed on ABI 3730XL DNA sequencer using the BigDye Terminator v1.1 Cycle Sequencing Kit. The sequencing results were compared with the reference sequence GenBank NM_014712.2 (*SETD1A* cDNA). This process is according to previously published procedures[1].

### Fly Stocks

The following fly stocks were obtained from the Bloomington Drosophila Stock Center: *mCherry*$^{RNAi}$ (stock #35785), *Set1*$^{RNAi1}$ (stock #33704), *Set1*$^{RNAi2}$ (stock #38368), *Set1*$^{RNAi3}$ (stock #40931), *R14H06-Gal4* (stock #48667), and *actin-Gal4* (stock #25374).

### Procedures Courtship Conditioning

Newly enclosed control (genotype: *mCherry*$^{RNAi}$*/R14H06-Gal4*) or knockdown (genotypes: *Set1*$^{RNAi1}$*/R14H06-Gal4*, *Set1*$^{RNAi2}$*/R14H06-Gal4*, *Set1*$^{RNAi3}$*/R14H06-Gal4*) males were isolated for 4 days. Individual males were then trained by introducing a single premated female into a training chamber. During the training period premated females reject male courtship attempts. This causes the male to learn, resulting in suppression of futile courtship behavior in subsequent pairings. For short-term memory (STM), males were trained for 1 hour and then re-isolated for 1 hour before testing courtship behavior with a new premated female. For long-term memory (LTM), we used an 8 hour training period, followed by a 20-24 hour rest before testing. For each fly pair a courtship index (CI) was calculated, which is the proportion of time spent courting over 10 minutes. CIs are presented in boxplots to show the distribution of the raw data. To test for memory, the mean CI of naïve flies was compared to that of the trained flies using a Mann-Whitney test (GraphPad Prism version 7.03). A significant reduction in mean CI from naïve to trained flies suggests normal memory. The learning index (LI) represents the percentage of reduction in courtship behavior due to training and is used to directly compare memory ability between different genotypes. LI is a single value that is calculated from the CIs using the formula: LI = (mean-$CI_{naïve}$ – mean-$CI_{trained}$)/ mean-$CI_{naïve}$. A randomization test[2] was used for statistical comparison of LIs between genotypes and was performed with a custom bootstrapping R script[3,4].
