## Supplementary Data 2 Results for "Characterization of *SETD1A* haploinsufficiency in humans and *Drosophila* defines a novel neurodevelopmental syndrome"

### Growth

In the majority of cases (9/14) the pregnancy was uneventful, while in three cases, a history of miscarriages was reported. In cases 13 (dizygotic twin pregnancy) and 14, the pregnancy was complicated by intrauterine growth retardation. In another case (10), oligohydramnion was reported. A low penetrance of growth abnormalities was observed within our cohort, including macrocephaly (3/13), both tall (3/15) and short (1/15) stature, and overweight (3/15).

### Miscellaneous

Skin abnormalities are reported in 5/15 individuals. One individual (case 9) had hemangioma, where hemangiomas have previously been reported in 2 cases[1]. Recurrent infections of the ears and respiratory tract were seen in 7/15 individuals. Other reported clinical features of importance are visual (6/15) and hearing (2/15) impairments, including one case of sensorineural hearing impairment and one case of conductive hearing impairment.

### Further evaluation of highlighted cases

#### *Case 5*

Case 5 had major immune and pulmonary problems, including common variable immune deficiency, decreased blood antibody levels, pulmonary hypertension and chronic interstitial pulmonary disease. As a result, he was in constant need of respiratory support, and died at the age of 4 years and 4 months from respiratory complications. The rest of the patients in our cohort and in the previously reported patients did not experience immune nor pulmonary problems, therefore it is unlikely that this patient's *SETD1A* variant attributed to the described medical condition. Taken together, further studies identifying novel cases with *SETD1A* Loss-of-Function variants are needed to reveal if this more severe phenotype is associated with the syndrome.

#### *Case 8*

This patient has a copy number variation (CNV): namely 16p13.11dup. Unfortunately this CNV has not been analysed in the parents. 16p13.11dup is associated with behavioral abnormalities and certain skeletal manifestations. Behavioral abnormalities include attention deficit/hyperactivity disorder, aggression and disruptive temperament, and autistic spectrum disorders. Skeletal manifestations include hypermobility, craniosynostosis and polydactyly[2].

This patient is also carrier of a maternally inherited *RBBP8* variant (c.1931del). *RBBP8* is known to cause Seckel Syndrome. Seckel Syndrome is a recessive disorder characterized by growth retardation, microcephaly with mental retardation, and a distinct facial appearance[3]. Since the patient only has a heterozygous variant, and there is no question of growth retardation or microcephaly (height is +0.7 SD, head circumference -0.3 SD), this additional variant seems unlikely to contribute to the phenotype.

#### *Case 12*

This individual has an additional, not de novo (maternally inherited) heterozygous *CNTNAP2* variant. Homozygous (or compound heterozygous) *CNTNAP2* variants are associated with Pitt-Hopkins-like syndrome-1[4].

#### *Case 13*

Case 13 presented with a missense variant (c.4495T>G, p.Tyr1499Asp). This variant lies in the COMPASS complex SET1 subunit, N-SET domain. We predict it to be disease causing (1) as the variant occurred *de novo*, (2) the patient showed features very similar to the majority of the cohort (i.e., developmental delay and distinctive facial features) (3) These patient cell lines displayed DNA damage repair defects that were comparable to previously observed RNAi-mediated depletion of *SETD1A*. Suggesting that this variant behaves as a Loss-of-Function (LoF) allele. and (4) multiple *in silico* prediction models indicated that this particular variant is pathogenic[5-8].

At birth, the boy presented with craniosynostosis, for which he received corrective surgery at the age of 1 year. In addition to the *SETD1A* missense variant, this patient also has a *de novo* variant in

*EFNB2*, a homologue of *EFNB1* which is known to be associated with craniosynostosis[9]. Though this is speculative, this feature might also be attributed to this variation in the *EFNB2* gene. Remarkably, craniosynostosis was also present in patient 3 of our *SETD1A* patient cohort. In case 3, no additional genetic variants were identified that could possibly be linked to craniosynostosis.

In addition, this patient has a third, maternally inherited variant c.3316C>T p.(His1106Tyr) in the *NHS* gene. Mutations in the *NHS* gene are associated with the Nance-Horan syndrome, or isolated cataracts[10,11]. Characteristics of the Nance-Horan syndrome itself include congenital cataracts, tooth abnormalities (abnormal shape of the teeth, screwdriver shape, absence of teeth, extra teeth or wide apart teeth) and approximately 30% of the boys with Nance-Horan syndrome have a developmental delay. The variants in *NHS* that lead to this syndrome are different than the variant described in case 13. The variants described in Nance-Horan syndrome mainly lead to a shortening of the NHS protein and thus a clear Loss-of-Function of the gene. The effect of the variant in this patient is not known. In addition, this patient has undergone several ophthalmological examinations, in which no abnormalities such as cataracts were found. Therefore it is our assessment that the variant in *NHS* is unlikely to be the cause of the described phenotype.
